# Effect of lysergic acid diethylamide (LSD) on reinforcement learning in humans

**DOI:** 10.1101/2020.12.04.412189

**Authors:** Jonathan W. Kanen, Qiang Luo, Mojtaba R. Kandroodi, Rudolf N. Cardinal, Trevor W. Robbins, Robin L. Carhart-Harris, Hanneke E.M. den Ouden

## Abstract

The non-selective serotonin 2A (5-HT_2A_) receptor agonist lysergic acid diethylamide (LSD) holds promise as a treatment for some psychiatric disorders. Psychedelic drugs such as LSD have been suggested to have therapeutic actions through their effects on learning. The behavioural effects of LSD in humans, however, remain largely unexplored. Here we examined how LSD affects probabilistic reversal learning in healthy humans. Healthy volunteers received intravenous LSD (75μg in 10 mL saline) or placebo (10mL saline) in a within-subjects design and completed a probabilistic reversal learning task. Participants had to learn through trial and error which of three stimuli was rewarded most of the time, and these contingencies switched in a reversal phase. Computational models of reinforcement learning were fitted to the behavioural data to assess how LSD affected the updating (“learning rates”) and deployment (“reinforcement sensitivity”) of value representations during choice, as well as “stimulus stickiness”, which assays choice repetition irrespective of reinforcement history. Conventional measures assessing sensitivity to immediate feedback (“win-stay” and “lose-shift” probabilities) were unaffected, whereas LSD increased the impact of the strength of initial learning on perseveration. Computational modelling revealed that the most pronounced effect of LSD was enhancement of the reward learning rate. The punishment learning rate was also elevated. Stimulus stickiness was decreased by LSD, reflecting heightened exploratory behaviour, while reinforcement sensitivity was unaffected. Increased reinforcement learning rates suggest LSD induced a state of heightened plasticity. These results indicate a potential mechanism through which revision of maladaptive associations could occur in the clinical application of LSD.

**Significance statement:** The psychedelic (“mind-manifesting”) drug LSD holds promise for the treatment of some psychiatric disorders. Theories have postulated its therapeutic potential centres on enhancing learning and flexible thinking. Here we provide substantiating empirical evidence by examining the computations underlying behaviour as healthy volunteers learned through trial and error under LSD. Viewing choice as based on representations of an action’s value, LSD increased the speed at which value was updated following feedback, which was more pronounced following reward than punishment. Behaviour was also more exploratory under LSD, irrespective of the outcome of actions. These results indicate that LSD impacted a fundamental belief-updating process inherent in the brain which can be leveraged to revise maladaptive associations characteristic of a range of mental disorders.

## Introduction

Research into lysergic acid diethylamide (LSD) as a potential therapeutic agent in psychiatry has been revitalised in recent years (Nutt and Carhart-Harris 2020; Vollenweider and Preller 2020). Theories on the putative beneficial effects of LSD on mental health centre on its effects on learning and plasticity (Carhart-Harris and Nutt 2017), yet few studies have examined its effect on human behaviour. LSD acts principally but not exclusively as an agonist at the serotonin (5-HT; 5-hydroxytryptamine) 2A [5-HT_2A_] receptor (Marona-Lewicka et al. 2005, 2007; Nichols 2016). Indeed, blocking 5-HT_2A_ receptors inhibits the psychedelic effects of LSD (Nichols 2016). The 5-HT_2A_ receptor is involved in plasticity (Barre et al. 2016; Vaidya et al. 1997) and its modulation represents a putative neurobiological mechanism through which LSD could facilitate the revision of maladaptive associations (Carhart-Harris and Nutt 2017). Indeed, LSD and 5-HT_2A_ agonists have been shown to improve associative learning in non-human animals (Harvey 2003; Harvey et al. 1988; Romano et al. 2010; Schindler et al. 1986). However, studies of human learning and cognitive flexibility under the influence of psychedelic drugs using objective tests (rather than subjective experience) are limited in number (Pokorny et al. 2019). Here we tested whether LSD altered probabilistic reversal learning in healthy volunteers, and explored how LSD altered underlying learning mechanisms, using reinforcement learning models.

Serotonin is critically involved in adapting behaviour flexibly as environmental circumstances change (Barlow et al. 2015; Brigman et al. 2010; Clarke et al. 2004; Matias et al. 2017; Rygula et al. 2015), as well as processing aversive outcomes (Bari et al. 2010; Chamberlain et al. 2006; Cools et al. 2008; Crockett et al. 2009; Dayan and Huys 2009; Deakin 2013; den Ouden et al. 2013; Geurts et al. 2013). Both can be modelled in a laboratory setting using probabilistic reversal learning (PRL) paradigms. In these, individuals learn by trial and error the most adaptive action, in an “acquisition” stage, and this rule eventually changes in a “reversal” phase (Lawrence et al. 1999). Profound neurotoxin-induced depletion of serotonin from the marmoset orbitofrontal cortex (OFC) causes perseverative, stimulus-bound behaviour (Walker et al. 2009) – an impaired ability to update action upon reversal (Clarke et al. 2004). At the same time, acute administration of selective serotonin reuptake inhibitors (SSRIs), which can paradoxically lower serotonin concentration (Nord et al. 2013), has resulted in an increased sensitivity to negative feedback (referred to as “lose-shift” behaviour) in healthy humans (Chamberlain et al. 2006; Skandali et al. 2018) and rats (Bari et al. 2010).

In addition to affecting the serotonin system, LSD has dopamine type 2 (D2) receptor agonist properties (Marona-Lewicka et al. 2005, 2007; Nichols 2004). Dopamine is particularly well known to play a fundamental role in learning from feedback (Schultz 2019; Schultz et al. 1997) putatively mediating plasticity changes during associative learning (Shen et al. 2008; Yin and Knowlton 2006). Meanwhile, dopamine depletion of the marmoset caudate nucleus, like serotonergic OFC depletion, also induced perseveration (Clarke et al. 2011). Additionally, there is a body of evidence, across species, that D2-modulating agents affect instrumental reversal learning (Boulougouris et al. 2009; Kanen et al. 2019; Lee et al. 2007).

The aim of the current study was to examine the effects of LSD on learning in humans, to inform the psychological mechanisms by which LSD could have salubrious effects on mental health. To do so, we tested the acute effects of LSD on PRL, in a placebo-controlled study of healthy human volunteers. We predicted LSD modulates either sensitivity to negative feedback or the impact of learned values on subsequent perseverative behaviour (den Ouden et al. 2013). Measuring “staying” (repeating a choice) or “shifting” (choosing another stimulus) after wins or losses assesses sensitivity to immediate reinforcement, but does not account for the integration of feedback history across multiple experiences to influence behaviour (Daw 2011). We applied computational models of reinforcement learning to test the hypothesis that LSD alters the rate at which value is updated following reward or punishment. Through modelling we additionally investigated whether LSD affects the degree to which behaviour is stimulus-driven (“stimulus sticky”), independent of an action’s outcome.

## Methods and Materials

### Subjects and drug administration

Nineteen healthy volunteers, over the age of 21, attended two sessions at least two weeks apart where they received either intravenous LSD (75μg in 10 mL saline) or placebo (10mL saline), in a single-blind within-subjects balanced-order design. All participants provided written informed consent after briefing on the study and screening. Participants had no personal history of diagnosed psychiatric disorder, or immediate family history of a psychotic disorder. Other inclusion criteria were normal electrocardiogram (ECG), routine blood tests, negative urine test for pregnancy and recent recreational drug use, negative breathalyser test for recent alcohol use, alcohol use limited to less than 40 units per week, and absence of a significant medical condition. Participants had previous experience with a classic psychedelic drug (e.g. LSD, mescaline, psilocybin/magic mushrooms, or DMT/ayahuasca) without an adverse reaction, and had not used these within six weeks of the study. Screening was conducted at the Imperial College London Clinical Research Facility (ICRF) at the Hammersmith Hospital campus, and the study was carried out at the Cardiff University Brain Research Imaging Centre (CUBRIC). Participants were blinded to the condition but the experimenters were not. A cannula was inserted and secured in the antecubital fossa and injection was performed over the course of two minutes. Participants reported noticing subjective effects of LSD 5 to 15 minutes after dosing. The PRL task was administered approximately five hours after injection. Once the subjective drug effects subsided, a psychiatrist assessed suitability for discharge. This experiment was part of a larger study, the data from which are published elsewhere (e.g. Carhart-Harris et al. 2016). Additional information, including subjective ratings, can be found in Carhart-Harris et al. (2016).

### Probabilistic reversal learning task

A schematic of the task is shown in Figure 1A. On every trial, participants could choose from three visual stimuli, presented at three of four randomised locations on a computer screen. In the first half of the task (40 trials), choosing one of the stimuli resulted in positive feedback in the form of a green smiling face on 75% of trials. A second stimulus resulted in positive feedback 50% of the time, whilst the third stimulus yielded positive feedback on only 25% of trials. Negative feedback was provided in the form of a red frowning face. The first stimulus that was selected, was defined as the initially rewarded stimulus; the choice on trial 1 always resulted in reward. The second stimulus that was selected was defined as the mostly punished stimulus, and by definition the 3^rd^ stimulus was then the “neutral” stimulus. After 40 trials, the most and least optimal stimuli reversed, such that the stimulus that initially was correct 75% of the time was then only correct 25% of the time, and likewise the 25% correct stimulus then resulted in positive feedback on 75% of trials. This is a novel version (Kandroodi et al. 2020) of a widely used PRL task (Lawrence et al. 1999; den Ouden et al. 2013): novel due to the addition of a 50% “neutral” stimulus in order to distinguish learning to select the mostly rewarding stimulus from learning to avoid the mostly punishing stimulus.

**Figure 1.**
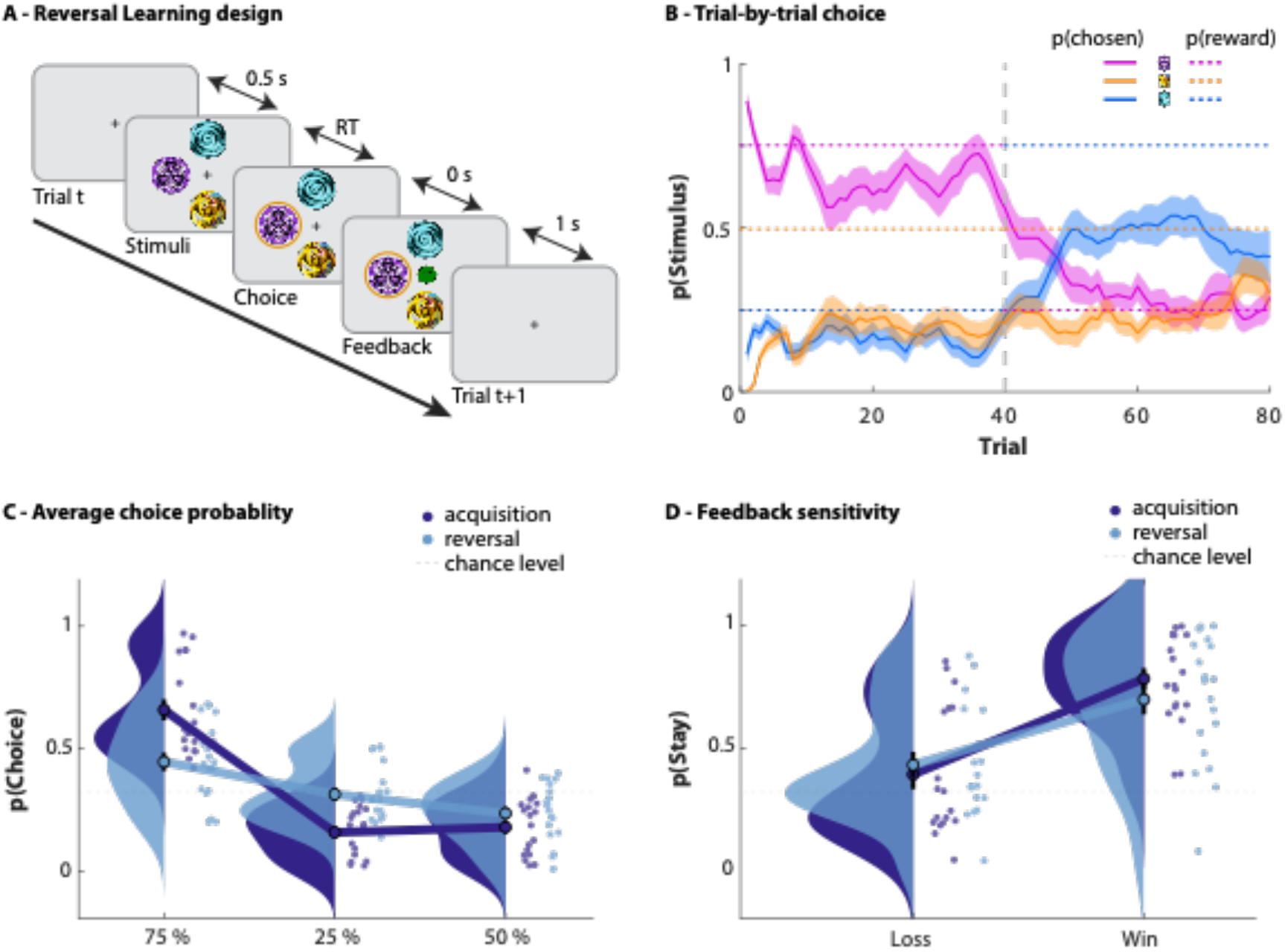
**A)** Schematic of the probabilistic reversal learning task. Subjects chose one of three stimuli. The timeline of a trial is depicted: stimuli appear, a choice is made, the outcome is shown, a fixation cross is presented during the intertrial interval, stimuli appear for the next trial (etc.) (RT, reaction time). One stimulus delivered positive feedback (green smiling face) with a 75% probability, one with 50%, and one with 25%. The probabilistic alternative was negative feedback (red sad face). Midway through the task, the contingencies for the best and worst stimuli swapped. s = seconds. **B)** Trial-by-trial average probability of choosing each stimulus, averaged over subjects and sessions, collapsed across LSD and placebo sessions. A sliding 5-trial window was used for smoothing. The vertical dotted line indicates the reversal of contingencies. Shading indicates 1 standard error of the mean (SE). **C)** Distributions depicting the average per-subject probability (scattered dots) of choosing each stimulus during the acquisition (shown in dark blue) and reversal (light blue) phases, collapsed across LSD and placebo sessions. Mean value for each distribution is illustrated with a single dot at the base of each distribution, and the mean values for the probability of choosing different stimuli in each phase are connected by a line. One SE is shown by black error bars around the mean value. Horizontal dotted line indicates chance-level stay-behaviour (33%). **D)** Distributions depicting the average per-subject probability (scattered dots) of repeating a choice (staying) after receiving positive or negative feedback during the acquisition (dark blue) and reversal (light blue) phases, collapsed across LSD and placebo sessions. Horizontal dotted line indicates chance-level stay-behaviour (33%).

### Conventional analysis of behaviour

We examined whether LSD impaired participants’ overall ability to perform the task by analysing the number of responses made to each stimulus during the acquisition and reversal phases. We measured feedback sensitivity by determining whether participants stayed with the same choice following positive or negative feedback (win-stay or lose-stay). The win-stay probability was defined as the number of times an individual repeated a choice after a win, divided by the number of trials on which positive feedback occurred (opportunities to stay after a win). Lose-stay probability was calculated in the same manner: number of times a choice was repeated following a loss, divided by the total losses experienced. Note that in previous studies with a choice between only two stimuli (or responses), this metric is usually referred to as “win-stay / lose-shift”, which also captures the tendency to repeat (rather than switch) responses following a win, and the tendency to switch (rather than repeat) choices following a loss. Random choice would result in 50% win-stay and 50% lose-shift; however, in the current paradigm with 3 stimuli, this base rate is 33% (win-)stay and 67% (lose-)shift. We therefore encode both variables with respect to the stay (rather than shift) rate, but they are still conceptually identical to earlier studies. Perseveration was defined according to den Ouden et al. (2013) and was assessed based on responses in the reversal phase. A perseverative error occurred when two or more (now incorrect) responses were made to the previously correct stimulus, and these errors could occur at any point in the reversal phase. The first trial in the reversal phase (trial 41 of 80) was excluded from the perseveration analysis, however, as at that point behaviour cannot yet be shaped by the new feedback structure. Note again that this metric is not entirely identical to the previous studies cited employing two stimuli, as the base-rate choice for each stimulus is now 1/3, so the “chance” level of perseverative errors is lower. Null hypothesis significance tests used *α* = 0.05.

### Computational modelling of behaviour

#### Model fitting, comparison, and interpretation

These methods are based on our previous work (Kanen et al. 2019). We fitted three reinforcement learning (RL) models to the behavioural data using a hierarchical Bayesian method, via Hamiltonian Markov chain Monte Carlo sampling implemented in Stan 2.17.2 (Carpenter et al. 2017). Convergence was checked according to 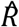, the potential scale reduction factor measure (Gelman et al. 2012; Brooks and Gelman 1998), which approaches 1 for perfect convergence. Values below 1.2 are typically used as a guideline for determining model convergence (Brooks and Gelman 1988). We assumed the three models had the same prior probability (0.33). Models were compared via a bridge sampling estimate of the marginal likelihood (Gronau et al. 2017a), using the “bridgesampling” package in R (Gronau et al. 2017b). Bridge sampling directly estimates the marginal likelihood, and therefore the posterior probability of each model given the data (and prior model probabilities), as well as the assumption that the models represent the entire group of those to be considered. Posterior distributions were interpreted using the 95% highest posterior density interval (HDI), which is the Bayesian “credible interval”. Parameter recovery for this modelling approach has been confirmed in a previous study (Kanen et al. 2019).

The Bayesian hierarchy consisted of “drug condition” at the highest level, and “subject” at the level below. For each parameter, each drug condition (e.g. LSD) had its own mean (with a prior that was the same across conditions, i.e. with priors that were unbiased with respect to LSD versus placebo). This was then merged with the intersubject variability (assumed to be normally distributed; mean 0 by definition, standard deviation determined by a further prior). The priors used for each parameter are shown in Table 1. For instance, the learning rate for a given subject under LSD was taken as: the group mean LSD value for learning rate, plus the subject-specific component of learning rate. The learning rate for a given subject under placebo was taken as: the group mean placebo value for learning rate, plus the subject-specific component of learning rate for the same subject. This method accounts for the within-subjects structure of the study design. This was done similarly (and separately) for all other model parameters.

**Table 1.** Prior distributions for model parameters

| Model parameters | Models using each parameter | Range | Prior | Reference |
| --- | --- | --- | --- | --- |
| reward learning rate, $\alpha^{rew}$ | 1, 3 | [0, 1] | Beta(1.2, 1.2) | den Ouden et al. (2013) |
| punishment learning rate, $\alpha^{pun}$ | 1, 3 | [0, 1] | Beta(1.2, 1.2) | den Ouden et al. (2013) |
| combined reward/punishment learning rate, $\alpha^{reinf}$ | 2 | [0, 1] | Beta(1.2, 1.2) | den Ouden et al. (2013) |
| reinforcement sensitivity, $\tau^{reinf}$ | 1, 2, 3 | [0, $+\infty$] | Gamma($\alpha=4.82$, $\beta=0.88$) | Gershman (2016) |
| stimulus stickiness, $\tau^{stim}$ | 2, 3 | $[-\infty, +\infty]$ | Normal(0, 1) | Christakou et al. (2013) |
| **Intersubject variability in parameters** |  |  |  |  |
| Intersubject standard deviations for $\alpha^{rew}$, $\alpha^{pun}$, $\alpha^{reinf}$, $\tau^{loc}$ | As above | [0, $+\infty$] | Half-normal: Normal(0, 0.05) constrained to $\geq 0$ | Kanen et al. (2019) |
| Intersubject standard deviations for $\tau^{reinf}$ | As above | [0, $+\infty$] | Half-normal: Normal(0, 1) constrained to $\geq 0$ | Kanen et al. (2019) |

*rew* reward, *pun* punishment, *reinf* reinforcement, *stim* stimulus

To determine the change (LSD – placebo) in parameters, we calculated [group mean LSD learning rate] – [group mean placebo learning rate] for each of the ~8,000 simulation runs and tested them against zero via the HDI. This approach also removes distributional assumptions and provides an automatic multiple comparisons correction (Gelman and Tuerlinckx 2000; Gelman et al. 2012; Kruschke 2011).

#### Models

The parameters contained in each model are summarised in Tables 1 and 2. With Model 1, we tested the hypothesis that positive versus negative feedback guides behaviour differentially, and that LSD affects this. We augmented a basic RL model (Rescorla & Wagner 1972) with separate learning rates for reward *α^rew^* and punishment *α^pun^*. Positive feedback led to an increase in the value *V_i_* of the stimulus *i* that was chosen, at a speed governed by the *reward learning rate α^rew^*, via *V*_*i,t*+1_ ← *V_i,t_* + *α^rew^*(*R_t_* – *V_i,t_*). *R_t_* represents the outcome on trial *t* (defined as 1 on trials where positive feedback occurred), and (*R_t_* – *V_i,t_*) the prediction error. On trials where negative feedback occurred, *R_t_* = 0, which led to a decrease in value of *V_i_* at a speed governed by the *punishment learning rate α^pun^*, according to *V*_*i,t*+1_ ← *V_i,t_* + *α^pun^*(*R_t_* – *V_i,t_*). Stimulus value was incorporated into the final quantity controlling choice according to *Q^reinf^_t_* = *τ^reinf^V_t_*. The additional parameter *τ^reinf^*, termed *reinforcement sensitivity*, governs the degree to which behaviour is driven by reinforcement history. The quantities *Q* associated with the three available choices, for a given trial, were then input to a standard softmax choice function to compute the probability of each choice:

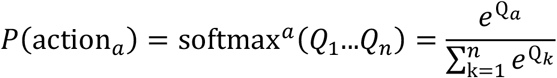

for *n* = 3 choice options. The probability values for each trial emerging from the softmax function (the probability of choosing stimulus 1) were fitted to the subject’s actual choices (did the subject choose stimulus 1?). Softmax inverse temperature was set to *β* = 1, and as a result the reinforcement sensitivity parameter (*τ^reinf^*) directly represented the weight given to the exponents in the softmax function.

**Table 2.** Model comparison

| Rank | Name | Parameters | log marginal likelihood | log posterior P(model) |
| --- | --- | --- | --- | --- |
| 2 | Model 1 | $\alpha^{rew}$, $\alpha^{pun}$, $\tau^{reinf}$ | -2401.49 | -33.28 |
| 3 | Model 2 | $\alpha^{reinf}$, $\tau^{reinf}$, $\tau^{stim}$ | -2428.52 | -60.32 |
| 1 | Model 3 | $\alpha^{rew}$, $\alpha^{pun}$, $\tau^{reinf}$, $\tau^{stim}$ | -2368.21 | 0 |

*rew* reward, *pun* punishment, *reinf* reinforcement, *stim* stimulus

Model 2 again augmented a simple RL model, but now also described the tendency to repeat a response, irrespective of the outcome that followed it (in other words, the tendency to “stay” regardless of outcome). With Model 2 we tested the hypothesis that LSD affects this basic perseverative tendency. This was implemented using a “stimulus stickiness” parameter *τ^stim^*. The stimulus stickiness effect was modelled as *Q^stim^_t_* = *τ^stim^s_t−1_*, where *s_t−1_* was 1 for the stimulus that was chosen on the previous trial and was 0 for the other two stimuli. In this model we used only a single learning rate *α^reinf^*. Positive reinforcement led to an increase in the value *V_i_* of the stimulus *i* that was chosen, at a speed controlled by the *learning rate α^reinf^*, via *V*_*i,t*+1_ ← *V_i,t_* + *α^reinf^*(*R_t_* – *V_i,t_*). The final quantity controlling choice incorporated the additional stickiness parameter as *Q_t_* = *Q^reinf^_t_* + *Q^stim^_t_*. Quantities *Q*, corresponding to the three choice options on a given trial, were then fed into the softmax function as above. It should be noted that if *τ^stim^* is not in the model (or is zero), then *τ^reinf^* is mathematically identical to the notion of softmax inverse temperature typically implemented as *β*. The notation *τ^reinf^* is used, however, because it contributes to *Q^reinf^_t_* but not to *Q^stim^_t_*. A standard implementation of *β*, by contrast, would govern the effects of both *Q^reinf^_t_* and *Q^stim^_t_* by weighting the sum of the two (*Q_t_*).

Model 3 was the full model that incorporated separate reward and punishment learning rates as well as the stimulus stickiness parameter. With Model 3, we tested the hypothesis that LSD affects both how positive versus negative feedback guides behaviour differentially, and how LSD affects a basic perseverative tendency. Again, the final quantity controlling choice was determined by *Q_t_* = *Q^reinf^_t_* + *Q^stim^_t_*.

## Results

### Learning and perseveration

First, we verified that LSD did not impair participants’ overall ability to perform the task. Behavioural performance is depicted in Figure 1 and 2. To examine whether LSD affected the number of times each stimulus was chosen, repeated-measures analysis of variance (ANOVA) was conducted with drug (LSD, placebo), phase (acquisition, reversal), and stimulus type (75%, 50%, or 25% rewarded) as within-subjects factors. This revealed a main effect of stimulus (*F*_1,23_ = 30.66, *p* = 3 × 10^-6^, *η_p_^2^* = .63), a stimulus × phase interaction (*F* = 28.62, *p* = 2 × 10^-6^, *η_p_^2^* = .61), and no interaction of LSD with stimulus or phase (*F*< 1.5, *p*> .24, *η_p_^2^*< .08, for terms involving LSD). The number of correct responses did not differ between placebo and LSD during the acquisition (paired-sample *t* test, *t*_18_ = 0.84, *p* = .4, *d* = .19) or reversal phases (*t*_18_ = 0.23, *p* = .8, *d* = .05).

**Figure 2.**
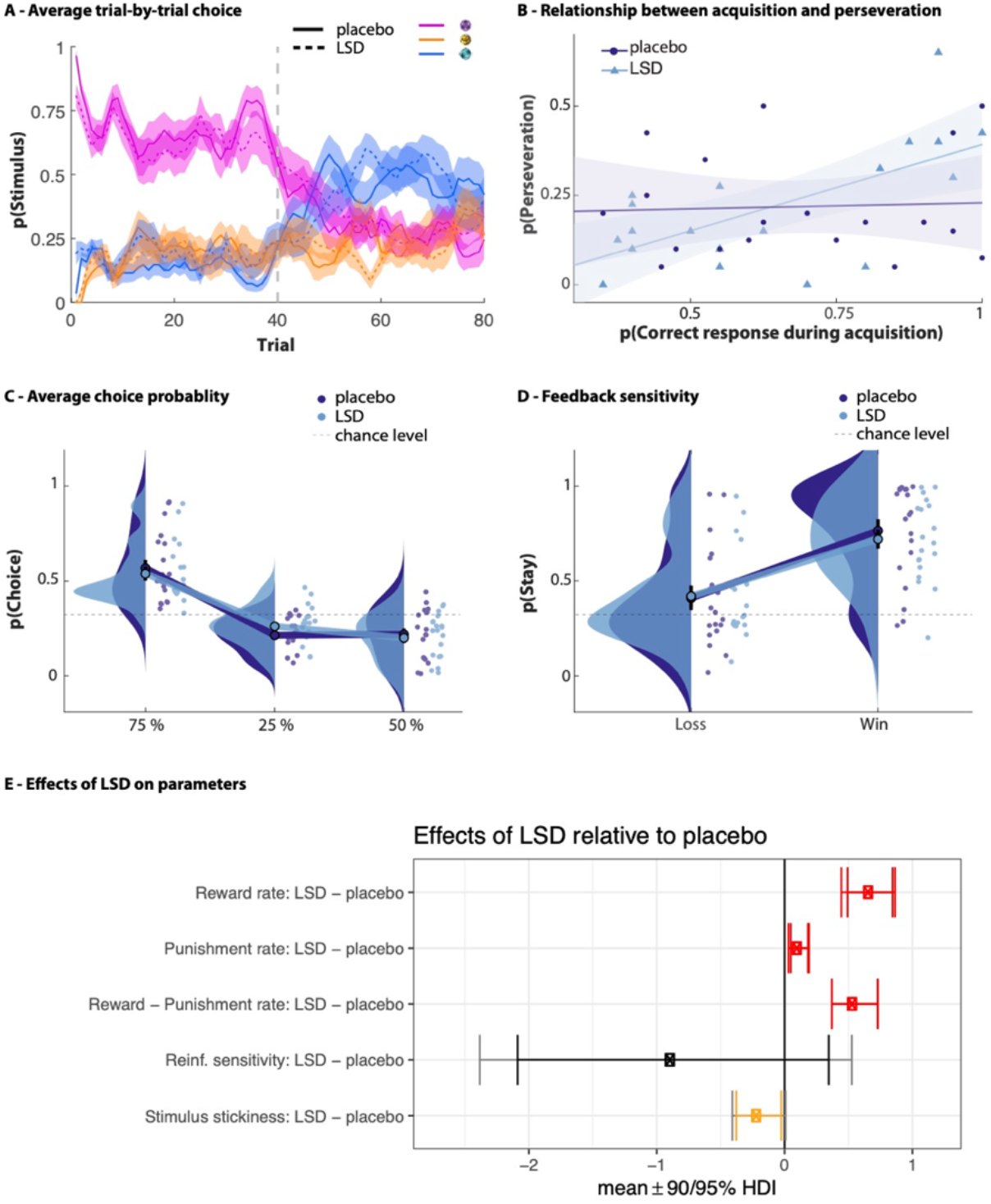
**A)** Trial-by-trial average probability of choosing each stimulus, averaged over subjects, separated by drug session. A sliding 5-trial window was used for smoothing. The vertical dotted line indicates the reversal of contingencies. Shading indicates 1 standard error of the mean (SE). **B)** Better initial learning was predictive of more perseveration on LSD and not on placebo. Shading indicates 1 SE. **C)** Distributions depicting the average per-subject probability (scattered dots) of choosing each stimulus while under placebo (shown in dark blue) and LSD (light blue). Mean value for each distribution is illustrated with a single dot at the base of each distribution, and the mean values for the probability of choosing different stimuli in each condition are connected by a line. One SE is shown by black error bars around the mean value. Horizontal dotted line indicates chance-level stay-behaviour (33%). D) Conventional analyses of feedback sensitivity were unaffected by LSD. Distributions depicting the average per-subject probability (scattered dots) of repeating a choice (staying) after receiving positive or negative feedback under placebo (dark blue) and LSD (light blue). Horizontal dotted line indicates chance-level stay-behaviour (33%). E) Effects of LSD relative to placebo on model parameters. Contrasts with the posterior 95% (or greater) highest posterior density interval [HDI] of the difference between means excluding zero (0 ∉ 95% HDI) are shown in red. Orange signifies 0 ∉ 90% HDI. The third row represents a difference of differences scores: [*α^rew^*_LSD_ – *α^pun^*_LSD_] – [*α^rew^*_placebo_ – *α^pun^*_placebo_].

We then examined the relationship between initial learning and perseveration, following den Ouden et al. (2013) (Figure 2B). LSD enhanced the relationship between the number of correct responses during the acquisition phase and the number of perseverative errors made during the subsequent reversal stage (acquisition correct responses [LSD minus placebo] versus reversal perseverative errors [LSD minus placebo]: linear regression coefficient *β* = .56, *p* = 0.002). Confirming this, making fewer errors during the acquisition phase predicted more perseverative errors when on LSD (*β* = 0.44, *p* = 0.003) but not when under placebo (*β* = 0.04, *p* = .8). Perseverative errors, a subset of all reversal errors, alone did not differ between conditions (*t*_18_ = 0.03, *p* = .98, *d* = .01).

### Feedback sensitivity

We next assessed whether LSD influenced individuals’ responses on trials immediately after positive versus negative feedback – whether participants stayed with the same choice after a win or a loss (win-stay/lose-stay; Figure 1D, 2D). Repeated-measures ANOVA with drug (LSD, placebo) and valence (win, loss) as within-subjects factors revealed a main effect of valence – participants “stayed” more after wins than losses (*F*_1,18_ = 37.76, *p* = 8.0 × 10^−6^, *η_p_^2^* = 0.68) – and no main effect of LSD (*F*_1,18_ = 0.20, *p* = .66, *η_p_^2^* = .01). There was also no interaction of valence × LSD (*F*_1,18_ = 0.63, *p* = .44, *η_p_^2^* = .03).

### Choice of reinforcement learning model

The core modelling results are displayed in Figure 2E. We fitted and compared three reinforcement learning models. Convergence was good with all three models having 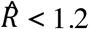. Behaviour was best characterised by a reinforcement learning model with four parameters (Table 2). The four parameters in the winning model were: 1) reward learning rate, which reflects the degree to which the chosen stimulus value is increased following a positive outcome (reward prediction error); 2) punishment learning rate, degree to which the chosen stimulus value is decreased following a negative outcome (punishment prediction error); 3) reinforcement sensitivity (comparable to inverse temperature), which is the degree to which the values learned through reinforcement contribute to final choice; and 4) “stimulus stickiness”, which indexes the tendency to get “stuck” to a stimulus and choose it because it was chosen on the previous trial, irrespective of outcome. The last two parameters resemble the explore/exploit trade-off: low values of stickiness or reinforcement sensitivity index two different types of exploratory behaviour.

### Reward and punishment learning rates

The reward learning rate was significantly elevated on LSD (mean 0.87) compared to placebo (mean 0.28) (with the posterior 99.9% highest posterior density interval [HDI] of the difference between these means excluding zero; 0 ∉ 99.9% HDI; Figure 2E). There was also an increased punishment learning rate under LSD (mean 0.48) relative to placebo (mean 0.39) (drug difference, 0 ∉ 99% HDI). Importantly, LSD increased the reward learning rate to a greater extent than the punishment learning rate ([*α^rew,LSD^* – *α^rew,placebo^*] – [*α^pun,LSD^* – *α^pun,placebo^*] > 0; drug difference, 0 ∉ 99% HDI).

### Stimulus stickiness and reinforcement sensitivity

Stimulus stickiness was lowered by LSD (mean 0.23) relative to placebo (mean 0.43) (drug difference, 0 ∉ 90% HDI; Figure 2E), which is a manifestation of increased exploratory behaviour. Reinforcement sensitivity was not modulated by LSD (LSD mean 4.70, placebo mean 5.57; no drug difference, 0 ∈ 95% HDI). This is in line with the absence of an effect of LSD on the tendency to ‘stay’ following reward or punishment (see conventional analysis above).

### Relationship between model parameters and conventional behavioural measures

Analyses to understand the relationship between computational and conventional measures were conducted, corrected for multiple comparisons, and are summarised in Supplementary Table 1. Given the initial finding on the relationship between better acquisition learning and perseveration, the first question addressed was whether the elevated reward learning rate from the computational model under LSD was predictive of the conventional measure of perseveration from den Ouden et al. (2013). Simple linear regression showed that under LSD, a higher reward learning rate predicted significantly more perseverative errors (*β* = 30.22, *p* = .02), whereas no such relationship was present when the same participants were under placebo (*β* = −.57, *p* = .95). Next we examined the relationship between the stimulus stickiness parameter from the computational model and the conventional measure of perseveration. Stimulus stickiness was not correlated with the conventional measure of perseveration, in either the placebo (*β* = 3.60, *p* = .43) or LSD (*β* = 7.67, *p* = .13) condition, indicating these were two independent processes. Further exploratory analyses are reported in Supplementary Table 1.

## Discussion

There has been a recent surge of interest in potential therapeutic effects of psychedelics, particularly LSD. Theorising on the mechanisms of such effects centres on their role in enhancing learning and plasticity. In the current study we tested these postulated effects of LSD in flexible learning in humans and find that LSD increased learning rates, exploratory behaviour, as well as the impact of previously learnt values on subsequent perseverative behaviour. Specifically, LSD increased the speed at which value representations were updated following prediction error (the mismatch between expectations and experience). Whilst LSD enhanced the impact of both positive and negative feedback, it augmented learning from reward significantly more than it augmented learning from punishment.

The observation that LSD enhanced learning rates may be particularly important for understanding the mechanisms through which LSD might be therapeutically useful. Psychedelic drugs have been hypothesised to destabilise pre-existing beliefs (i.e. relax prior beliefs or “priors”), making them amenable to revision (Carhart-Harris and Friston 2019). The notion of relaxed priors is directly compatible with increased reinforcement learning rates: in our study, LSD rendered subjects more sensitive to prediction errors, which naturally implies downweighting of prior beliefs (Carhart-Harris and Friston 2019). That LSD affected a fundamental belief-updating process is notable given that psychedelics are under investigation trans-diagnostically for diverse clinical phenomena including depression (Carhart-Harris et al. 2016, 2018; Ross et al. 2016), anxiety (Griffiths et al. 2016; Grob et al. 2011), alcohol (Bogenschutz et al. 2015) and nicotine abuse (Johnson et al. 2014), OCD (Moreno et al. 2006), and eating disorders (Lafrance et al. 2017); a unifying feature of these conditions is maladaptive associations in need of revision.

Behaviour was more exploratory overall under LSD irrespective of reinforcement history, reflected by lower estimates of the stimulus stickiness parameter. This is entirely consistent with theoretical accounts of psychedelic effects which have predicted increased exploratory tendencies (Carhart-Harris and Friston 2019). Another type of exploratory behaviour that is more commonly studied, low reinforcement sensitivity (a lower tendency to exploit a highly valued option), was unaffected by LSD: this highlights the utility of the stimulus stickiness parameter in parcellating and thus uncovering exploratory behaviour. That LSD lowered stimulus stickiness may also be clinically relevant: stimulus stickiness was recently shown to be abnormally high in cocaine and amphetamine use disorders, whilst reinforcement sensitivity was unaffected (Kanen et al. 2019).

Under LSD, better initial learning led to more perseverative responding. Importantly, perseveration (den Ouden et al. 2013) itself from the conventional analysis was not elevated by LSD nor did it correlate with stimulus stickiness (Supplementary Table 1), indicating these are two independent measures. The implication is that when a behaviour is newly and more strongly learned through positive reinforcement (i.e. the acquisition phase) under LSD, it may persist more strongly even when that action is no longer relevant (i.e. the reversal phase). This is orthogonal to an overall tendency towards exploration irrespective of reinforcement history (low stimulus stickiness, estimated from choice behaviour across both acquisition and reversal phases).

Given the broad effect of LSD on a range of neurotransmitter systems (Nichols 2004, 2016), it is not possible to determine the specific neurochemical mechanism underlying the observed LSD effects on learning. Nonetheless, obvious possibilities involve the serotonin and dopamine system, in particular 5-HT_2A_ and D2 receptors (Marona-Lewicka et al. 2005, 2007; Nichols 2004, 2016). Specifically, the psychological plasticity purportedly promoted by psychedelics is believed to be mediated through action at 5-HT_2A_ receptors (Carhart-Harris and Nutt 2017) via downstream enhancement of NMDA (N-methyl-D-aspartate) glutamate receptor transmission (Barre et al. 2016) and brain-derived neurotrophic factor (BDNF) expression (Vaidya et al. 1997). The hypothesis that the present reinforcement learning rate results are driven by serotonergic effects of LSD is supported by two recent studies in mice. Optogenetically stimulating dorsal raphé serotonin neurons enhanced reinforcement learning rates (Iigaya et al. 2018), whilst activation of these neurons tracked both reward and punishment prediction errors during reversal learning (Matias et al. 2017). Neurotoxic manipulation of serotonin in marmoset monkeys during PRL, meanwhile, altered stimulus stickiness (Rygula et al. 2015): this implicates a serotonergic mechanism underlying increased exploratory behaviour following LSD administration in the present study.

In addition to affecting the serotonin system, however, LSD also acts at dopamine receptors (Nichols 2004, 2016). Dopamine has long been known to play a crucial role in belief updating following reward (Schultz et al. 1997), and more recent evidence shows that dopaminergic manipulations may alter learning rates (Kanen et al. 2019; Schultz 2019; Swart et al. 2017). A dopaminergic effect would be in line with our previous study where genetic variation in the dopamine, but not serotonin transporter polymorphism, was associated with the same enhanced relationship between acquisition and perseveration as reported here under LSD (den Ouden et al. 2013).

Serotonin–dopamine interactions represent another candidate mechanism that could underlie the present findings. For example, stimulation of 5-HT_2A_ receptors in the prefrontal cortex of the rat, enhanced ventral tegmental area (VTA) dopaminergic activity (Bortolozzi et al. 2005). Indeed, the initial action of LSD at 5-HT_2A_ receptors has been proposed to sensitise dopamine neuron firing, which subsequently potentiates the direct dopaminergic effects of LSD (Nichols 2016). LSD action at D2 receptors, consequently, appears to be especially pronounced at a later time following LSD administration (Marona-Lewicka et al. 2005, 2007), which is relevant given the relatively long delay between LSD administration and performance of the current task (see Methods). However, arguing against a late dopaminergic effect is a previous study in rodents where the effects of LSD on reversal learning were consistent across four different time lags between drug administration and behavioural testing (King et al. 1974).

The result of enhanced coupling of acquisition learning and perseverative responding under LSD is in line with a recent study showing that LSD induced spatial working memory deficits and higher-order cognitive inflexibility in a set-shifting paradigm (Pokorny et al. 2019). Importantly, these effects were blocked by co-administration of the 5-HT_2A_ antagonist ketanserin (Pokorny et al. 2019), showing that the LSD-induced impairments were mediated by 5-HT_2A_ agonism, consistent with a 5-HT_2A_ mechanism underlying the present results.

LSD’s effects to increase acquisition-perseveration coupling and worsen set-shifting (Pokorny et al. 2019), in conjunction, suggest that what is newly learned through reinforcement under LSD is more “stamped in”, which may subsequently be harder to update. Whilst these findings are ostensibly at odds with the observation that LSD enhanced plasticity (through enhanced learning rates), these results can be reconciled by considering the timing of drug administration with respect to initial learning and tests of cognitive flexibility. In both the present experiment and the previous set-shifting study (Pokorny et al. 2019), all phases of learning (acquisition and reversal) were conducted after LSD administration. In contrast, when acquisition learning was conducted prior to LSD administration, LSD resulted in improved reversal learning (using a reversal paradigm in rats; King et al. 1974). Likewise, when acquisition learning was conducted prior to administration of a 5-HT_2A_ antagonist, reversal learning was impaired (Boulougouris et al. 2008). Collectively, these findings suggest that whether a prior belief is down- or upweighted under LSD may depend on whether the prior is formed before or during drug administration, respectively. This observation is of great relevance for a putative therapeutic setting, where maladaptive beliefs will have been formed before treatment.

In summary, the core result of this study was that LSD enhanced the rate at which humans updated their beliefs based on feedback. Learning rate was most enhanced by LSD when receiving reward, and to a lesser extent following punishment. LSD also increased exploratory behaviour, which was independent of reinforcement history. This study represents one of the few applications of objective measures to investigate fundamental cognitive processes in humans under LSD. These findings have implications for understanding the mechanisms through which LSD might be therapeutically useful for revising deleterious associations.

## Acknowledgements

This study was funded by the Walacea.com crowdfunding campaign and the Beckley Foundation, awarded to R.L.C-H. J.W.K. was supported by a Gates Cambridge Scholarship and an Angharad Dodds John Bursary in Mental Health and Neuropsychiatry, T.W.R. by a Wellcome Trust Senior Investigator Grant 104631/Z/14/Z, and H.E.M.d.O. by the Netherlands Organisation for Scientific Research, NWO. R.N.C.’s research is funded by the UK Medical Research Council (MC_PC_17213). Q.L. was supported by the National Key Research and Development Program of China (grant 2018YFC0910503), the National Natural Science Foundation of China (grant 81873909), the Natural Science Foundation of Shanghai (grant 20ZR1404900), the Shanghai Municipal Science and Technology Major Project (grant 2018SHZDZX01), and the Zhangjiang Lab.

## Supplementary information

### Simulation: Methods

We simulated behavioural data from the winning model to determine how behavioural patterns in the synthetic data compared to the raw data. Simulated data were analysed for win-stay probability, lose-stay probability, acquisition performance, and perseveration, as was done for the original raw data analysis. For each condition (placebo and LSD), we simulated 100 “virtual subjects” using the posterior mean parameters from that condition, from the winning model, per Kanen et al. (2019).

### Simulation: Results

Simulated behavioural data, generated using parameter estimates from the winning model, were analysed using conventional methods in order to assess whether the winning model could capture the observed effects of LSD on raw behaviour. Simulated data are shown in Supplementary Figure 1. Consistent with the original data, lose-stay probability was unaffected by LSD in the simulated behaviour (*t*_99_ = −0.37, p = .71, d = .03) and acquisition performance was also unaffected (*t*_99_ = 0.25, p = .81, d = .03). Perseveration was enhanced by LSD in the simulation (*t*_99_ = −2.24, *p* = 0.03, *d* = .22), which differs slightly from, yet is in line with, the original analyses showing an enhanced relationship between acquisition and perseveration under LSD. Linear regression examining whether correct responses during the acquisition phase (LSD minus placebo) predicted more perseverative errors in the reversal stage (LSD minus placebo) was not significant in the simulated data (*β* = 0.15, *p* = 0.13). Separate regressions for each condition also showed no significant relationship between acquisition performance and perseveration for LSD (*β* = 0.12, *p* = 0.25) or for placebo (*β* = 0.01, *p* = .91). Win-stay probability was diminished under LSD in the simulated data (*t*_99_ = 11.91, *p* = 8.21 × 10^−21^, *d* = 1.19) whereas it was unaffected by LSD in the raw data analysis.

**Supplementary Figure 1.**
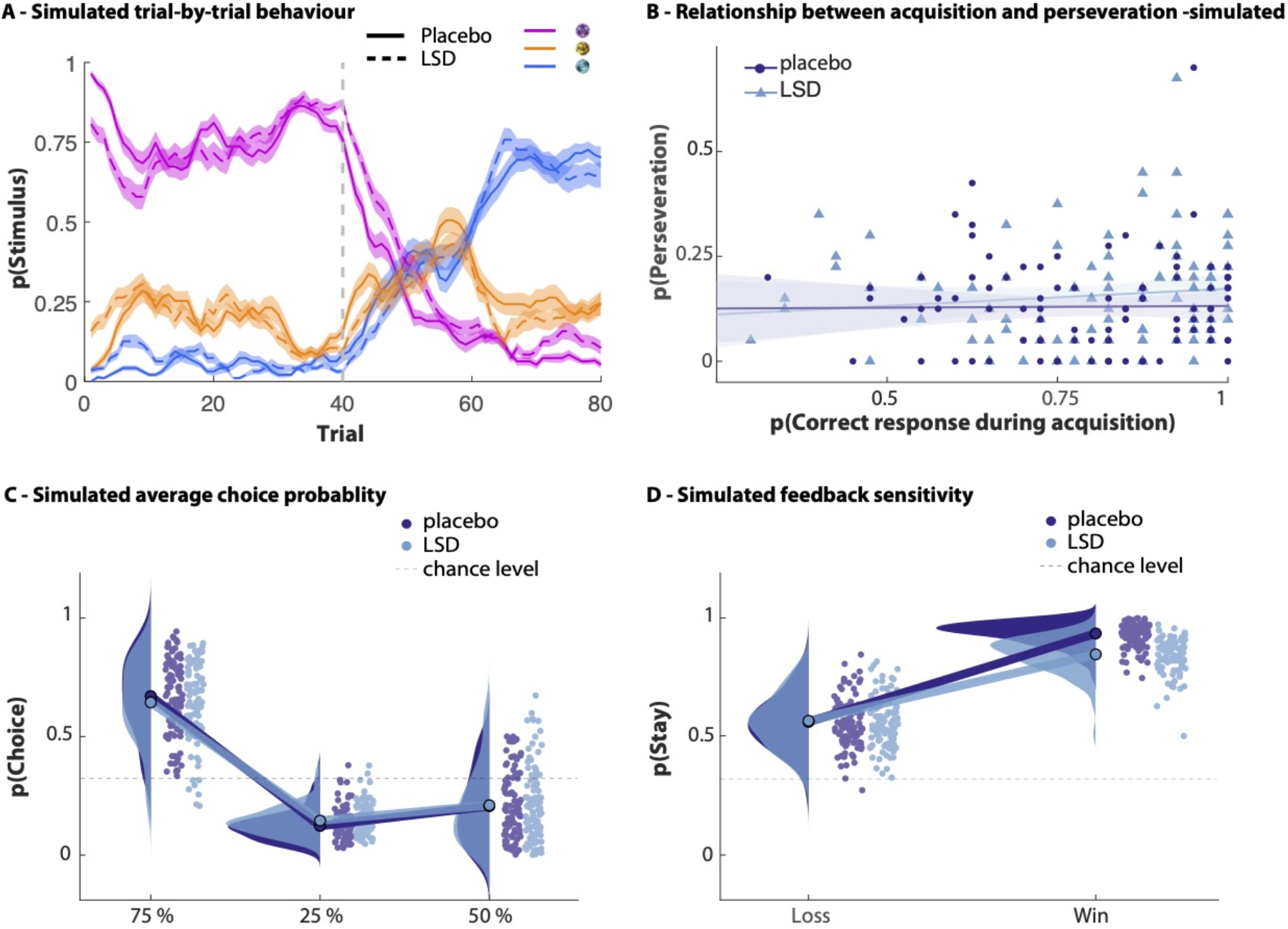
Simulated data. **A)** Trial-by-trial average probability of choosing each stimulus, averaged over simulated subjects, separated by drug session. A sliding 5-trial window was used for smoothing. The vertical dotted line indicates the reversal of contingencies. Shading indicates 1 standard error of the mean (SE). **B)** Relationship between initial learning and perseveration on LSD versus placebo in simulated data. Shading indicates 1 SE. **C)** Distributions depicting the average per-subject probability (scattered dots) of simulated subjects choosing each stimulus while under placebo (shown in dark blue) and LSD (light blue). Mean value for each distribution is illustrated with a single dot at the base of each distribution, and the mean values for the probability of choosing different stimuli in each condition are connected by a line. One SE is shown by black error bars around the mean value. Horizontal dotted line indicates chance-level stay-behaviour (33%). **D)** Distributions depicting the average per-subject probability (scattered dots) of simulated subjects repeating a choice (staying) after receiving positive or negative feedback under placebo (dark blue) and LSD (light blue). Horizontal dotted line indicates chance-level stay-behaviour (33%).

**Supplementary Table 1.** Summary of correlations between conventional behavioural measures and model parameters. ↑ significant positive correlation; ↓ significant negative correlation; – no significant correlation. All significant correlations survived correction for 32 comparisons, using the Benjamini-Hochberg method at q = .15 (Skandali et al. 2018).

|  | Acquisition Performance |  | Perseveration |  | Proportion Lose-Stay |  | Proportion Win-Stay |  |
| --- | --- | --- | --- | --- | --- | --- | --- | --- |
|  | Placebo | LSD | Placebo | LSD | Placebo | LSD | Placebo | LSD |
| **Reward Learning Rate, $\alpha^{rew}$** | –<br>$p = .86$ | –<br>$p = .053$ | –<br>$p = .95$ | ↑<br>$p = .02$ | –<br>$p = .79$ | –<br>$p = .37$ | –<br>$p = .55$ | –<br>$p = .11$ |
| **Punishment Learning Rate, $\alpha^{pun}$** | –<br>$p = .18$ | –<br>$p = .053$ | –<br>$p = .34$ | –<br>$p = .60$ | ↓<br>$p = .01$ | ↓<br>$p = .04$ | –<br>$p = .08$ | –<br>$p = .65$ |
| **Reinforcement Sensitivity, $\tau^{reinf}$** | ↑<br>$p = 1 \times 10^{-3}$ | ↑<br>$p = 3 \times 10^{-3}$ | –<br>$p = .26$ | ↑<br>$p = .02$ | ↑<br>$p = 1 \times 10^{-3}$ | ↑<br>$p = .01$ | ↑<br>$p = .01$ | ↑<br>$p = .02$ |
| **Stimulus Stickiness, $\tau^{stim}$** | ↑<br>$p = .02$ | –<br>$p = .16$ | –<br>$p = .43$ | –<br>$p = .13$ | ↑<br>$p = 2 \times 10^{-3}$ | ↑<br>$p = .03$ | –<br>$p = .07$ | ↑<br>$p = .02$ |

*rew* reward, *pun* punishment, *reinf* reinforcement, *stim* stimulus

